# Interspecific interactions regulate plant reproductive allometry in cereal-legume intercropping systems

**DOI:** 10.1101/2021.06.17.448782

**Authors:** Noémie Gaudio, Cyrille Violle, Xavier Gendre, Florian Fort, Rémi Mahmoud, Elise Pelzer, Safia Médiène, Henrik Hauggaard-Nielsen, Laurent Bedoussac, Catherine Bonnet, Guénaëlle Corre-Hellou, Antoine Couëdel, Philippe Hinsinger, Erik Steen Jensen, Etienne-Pascal Journet, Eric Justes, Bochra Kammoun, Isabelle Litrico, Nathalie Moutier, Christophe Naudin, Pierre Casadebaig

## Abstract

1. Calls for ecological principles in agriculture have gained momentum. Intercropping systems have long been designed by growing two, or more, annual crop species in the same field, aiming for a better resource use efficiency. However, assembly rules for their design are lacking. Notably, it is still unknown whether species performances are maximized during both the vegetative and reproductive phases given the sensitivity of reproductive allocation rules to resource limitation. Interestingly, ecological theory provides expectations regarding putative invariance of plant reproductive allometry (PRA) under non-limiting conditions for plant growth. Here we examined whether and how PRA changes in response to plant-plant interactions in intercropping systems, which can inform both ecological theory and the understanding of the functioning of intercropping systems.
2. We analyzed a dataset of 28 field cereal-legume intercropping trials from various climatic and management conditions across Western Europe. PRA was quantified in both mixing and single-species situations.
3. PRA was positively impacted in specific management conditions, leading to a greater increase in yield for a given increase in plant size. Variations in PRA were more beneficial for legumes grown in unfertilized mixture, which explains their use as a key component in actual intercrop systems. The response for cereals was similar but less pronounced in magnitude, and was greater under limiting resource conditions. Focusing on intercropping conditions, hierarchical competition (indicated by biomass difference between intercropped species) appears as a strong driver of the reproductive output of a given species.
4. Synthesis and applications. Plant reproductive allometry behaves in crop species in the same way as it does in wild species. However, contrary to theoretical expectations about an overall invariance of PRA, we highlighted taxon-specific and context-dependent effects of plant-plant interactions on PRA. This systematic deviation to PRA expectations could be leveraged to cultivate each species up to its reproductive optimum while accounting for the performance of the other, whether farmer’s objective is to favor one species or to reach an equilibrium in seed production. Sowing density and cultivar choice could regulate the biomass of each component, with specific targets derived from allometric relationships, aiming for an optimal reproductive allocation in mixtures.

## Introduction

Intercropping, i.e. combining at least two annual crop species in the same field for most part of their growing periods (Willey, 1979), is a promising way to move towards more sustainable agriculture (Lin, 2011; Li-li et al., 2015). Intercropped species are expected to use resources differently and more efficiently (e.g. Malézieux et al., 2009; Beillouin et al., 2019; Jensen et al., 2020). Many intercrops mix a cereal and a legume, with the underlying assumption that the cereal will benefit from the legume’s atmospheric nitrogen (N) fixation, thus decreasing the need for exogenous N fertilization (Thorsted et al., 2006; Malézieux, 2012; Gaba et al., 2018). The performance of intercropping systems has been studied from an agronomic perspective, focusing mainly on yield and N use (Hauggaard-Nielsen et al., 2009a; e.g. Bedoussac and Justes, 2010a; Naudin et al., 2010; Pelzer et al., 2012). While introducing species diversity into cropping systems could appear promising under low-input conditions, specific recommendations for the management of intercrops is in its infancy (Litrico and Violle, 2015). One reason for this is that the underlying mechanisms of the positive effect of the intercropping remain elusive, which makes it challenging to choose species and cultivars for these systems accurately. One key unsolved issue for identifying these mechanisms is how vegetative biomass translates into reproductive biomass and how reproductive allocation differs between sole cropping and intercropping situations. Bridging ecology and agronomy could help resolve this issue.

In ecology, the plant allometry literature has extensively analyzed the change in many key plant features as a function of size. Notably, a large body of theory indicates that plant reproductive output (grain yield for annual cropping systems) is a function of plant size (Weiner et al., 2009). It is based on metabolic optimization criteria, in which regulation processes and selection forces have similar influence on size-related traits across taxa (Enquist et al., 1999). It forms the basis of metabolic scaling theory (MST), which provides first principles of plant allometry laws (West et al., 1997, 1999). As a macroecological law, MST explains trait variation across several orders of magnitude of taxa, scales and body size. This body of theory attracts interest for the design and management of intercropping systems given the predictive power of universal scaling equations of MST (Deng et al., 2012).

The hypothesis of invariance at the origin of allometric scaling laws has been challenged. Poorter et al. (2015) highlighted that allometric scaling exponents differ among species. Vasseur et al. (2012) and Vasseur et al. (2018) highlighted variability in these exponents within the model species *Arabidopsis thaliana*, and demonstrated that this variability was genetically determined and environmentally regulated due to natural selection. Further, the influence of artificial selection on allometric constraints is not well understood due to the lack of comparisons of allometric relationships in crop species (Milla et al., 2015). The initial MST framework was designed along with plant observations in optimal conditions, i.e. where growth is not strongly limited by unfavorable abiotic or biotic conditions. Consequently, the influence of plant-plant interactions and soil resource limitations on deviations from MST expectations remains unknown (but see Coomes et al., 2011; Vasseur et al., 2018). Intercropping systems represent a unique opportunity to challenge allometric laws, in order to fine-tune them and assess the validity of their most basic assumptions. Understanding the influence of plant-plant interactions on reproductive strategies of intercropped species would improve the understanding, modeling and ultimately management of intercrops (Gaudio et al., 2019), particularly to drive each species to its potential reproductive output in relation to the other species and the cropping conditions. In this study, we analyzed how plant allometry is related to the performance of intercropped species and how this relationship is influenced by varying cropping conditions. Crop scientists and stakeholders, including farmers, are primarily interested in yield, often assessed in intercropping systems by the land equivalent ratio in order to calculate land-use efficiency (e.g. Yu et al., 2016). Finer analysis of intercrop performance would improve our understanding of the mechanisms underlying intercrop performance.

We examined the influence of plant-plant interactions on the allometric relationship between grain yield production and plant biomass in annual cereal-legume intercrops grown under a variety of climatic and cropping conditions in Western Europe, with the underlying objective to test the MST under non-optimal conditions, characterized here by the plant-plant interactions and soil nutrient limitations. Our analysis was based on 28 field experiments. The main objective of this study was to investigate how the reproductive allometric relationships of both plant families (cereals and legumes) changed depending on whether they were grown in a sole crop or with another crop. We also focused on the influence of N fertilization within each plant family and crop type (sole crop *vs.* intercrop). The strength of allometric relationships can indicate that the ratio of yield to plant size does not vary, as the relationship between these two variables is supposed to be invariant (Nee et al., 2005). This ratio is called “reproductive effort” in ecology (Cheplick, 2005) and “harvest index” in agronomy (Vega et al., 2000; Echarte and Andrade, 2003). It is often used to focus on allocation of biomass to reproductive organs and to differentiate performances of species and cultivars (Hay, 1995), which is a framework that is complementary to MST. Thus, we also assessed the influence of crop management on reproductive efforts of the two plant families.

## Materials and methods

### Field experiments

We collected a set of experiments that compared different species and cultivars under intercropping and sole-cropping conditions under a variety of management practices in 9 locations in five European countries (France, Denmark, Italy, Germany, and the United Kingdom) (Fig. 1). The experiments covered 28 environments (location × year), of which 15 were managed as organic farming and 13 as conventional farming, with a total of 34 intercropping situations (environment × species) and 62 sole-cropping situations. Since the experiments were not completely factorial, i.e. not all factors (cultivars, N fertilization, sowing density) were combined, we analyzed a total of 159 and 219 experimental units under intercropping and sole-cropping situations, respectively. In the experiments, 53% and 47% of the intercropped species were winter and spring crops, respectively. The mean temperature over the crop cycle (from sowing to harvest) ranged from 6.8-11.3 °C for winter crops and 12.3-15.1 °C for spring crops. Cumulative rainfall ranged from 278-713 mm for winter crops and 60-366 mm for spring crops.

**Fig. 1.**
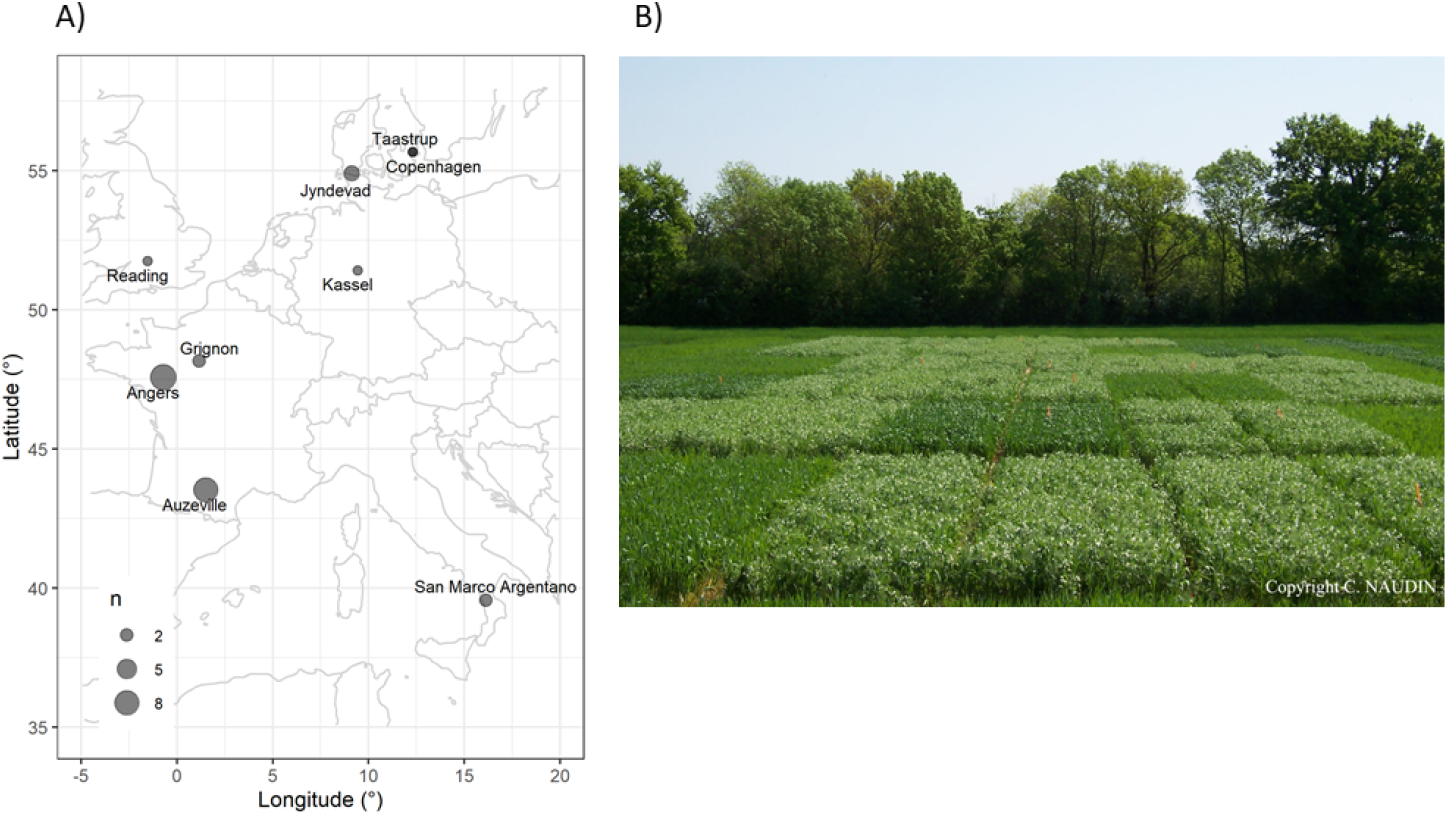
(A) Number of field experiments (size of the circle) conducted at each location and (B) example of a field experiment of winter wheat-pea intercrops (and their corresponding sole crops) conducted at the ARVALIS experimental station, near Angers, France (Source: C. Naudin).

Additional details on experimental designs and management practices are reported in Supplementary Material S1 and in the reference publications for 22 of the 28 experiments (Knudsen et al., 2004; Corre-Hellou et al., 2006; Hauggaard-Nielsen et al., 2008; Hauggaard-Nielsen et al., 2009a; b; Launay et al., 2009; Bedoussac and Justes, 2010b; a; Naudin et al., 2010, 2014; Pelzer et al., 2016; Tang et al., 2016). The set of experiments included annual cereal-grain legume intercrops and their corresponding sole crops, with i) barley (*Hordeum vulgare* L.), durum wheat (*Triticum turgidum* L.) and soft wheat (*Triticum aestivum* L.) as the cereals (only Poaceae), and ii) faba bean (*Vicia faba* L.) and pea (*Pisum sativum* L.) as the legumes (Fabaceae). The cross between crop species and cropping seasons resulted in five intercropping combinations: two spring intercrops (barley-faba bean and barley-pea) and three winter intercrops (durum wheat-faba bean, durum wheat-pea and soft wheat-pea). In all experiments, the two intercropped species were sown and harvested at the same time, with sowing dates ranging from March 11 to May 03 for spring crops and October 25 to December 15 for winter crops. Within a given cropping situation, variations were related mainly to i) the number of cultivars tested per crop species (ranging from 1-5); ii) the relative sowing density of each species (actual:reference sowing density ratio, 1.0 and 0.5 for sole crops and 0.3 - 0.7 for each of the two intercropped species); and iii) the N fertilization, with non-fertilized and N-fertilized situations, the latter ranging from 30-200 kg N.ha^−1^ (mean (± SD) = 95 ± 44 kg N.ha^−1^) (Table 1). To assess reproductive effort and allometry, all experiments measured at least three variables: grain yield (t.ha^−1^) at harvest, total aboveground biomass (t.ha^−1^, including grains, flowers, pods and ears) at maturity, and actual plant density (plant.m^−2^). Plant density was used to convert per-ha variables into per-capita variables (i.e. g.plant^−1^; Table 1).

**Table 1.**
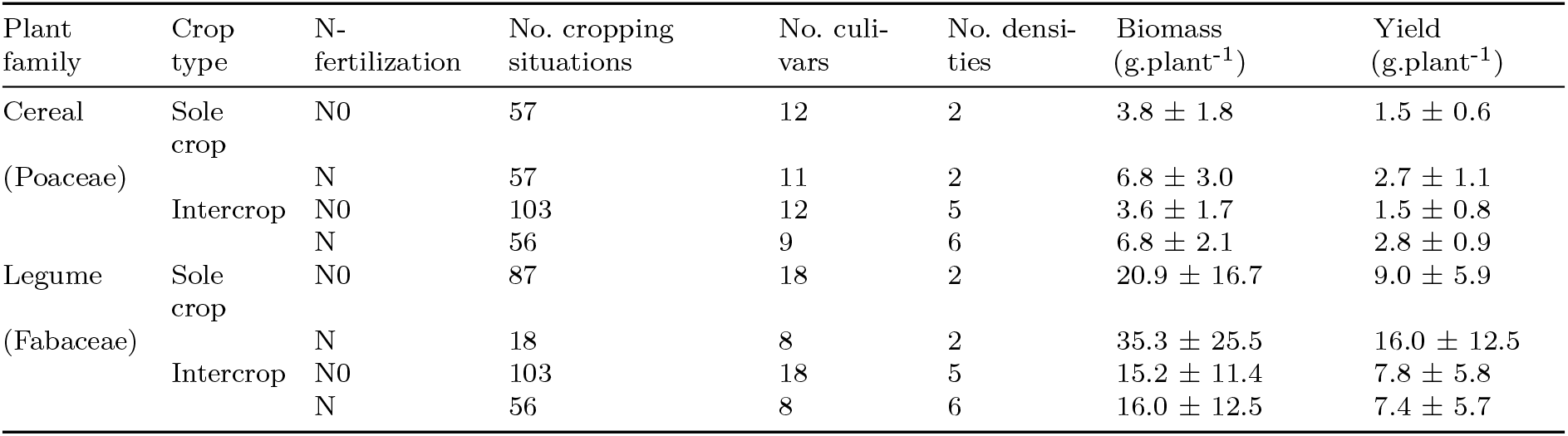
Cropping situations pooled in the database by plant family (Poaceae *vs.* Fabaceae), crop type (sole crop *vs.* intercrop) and nitrogen (N) fertilization (non-fertilized N0 *vs.* N-fertilized). Several cultivars and crop species densities (relative proportions) were represented for each factor combination (family × crop type × N fertilization). Mean (± SD) total plant aboveground dry biomass and grain yield were calculated for each factor combination.

### Data processing and analysis

Reproductive effort was calculated as the ratio of grain yield to total aboveground biomass at maturity, rather than final biomass, to avoid the influence of leaves that dropped before maturity (Unkovich et al., 2010). Analysis of variance (ANOVA) was performed using the *aov* function of the stats package of R software (R Core Team, 2019). When relevant (p < 0.05), means were separated using a Tukey or, when ANOVA assumptions were not met, Kruskal-Wallis test. We performed one-way ANOVAs within each plant family (Poaceae *vs.* Fabaceae) to test the influence of crop type (sole crop *vs.* intercrop) and N fertilization (non-fertilized *vs.* N-fertilized) on reproductive effort and its components (i.e. plant aboveground biomass and plant yield). The influence of N fertilization could not be assessed for legumes in sole crops due to unequal sample sizes (18 situations fertilized, 87 non-fertilized) (Table 1). For the same reason, differences between sole crops and intercrops for legumes could be assessed only under non-fertilized conditions.

We analyzed reproductive allometric relationships between plant grain yield and plant aboveground biomass thanks to standardized major axis analysis [SMA; Warton et al. (2006); Poorter and Sack (2012)] using the R *smatr* package (Warton et al., 2012), through the power relation *y* = *ax^b^*, where *y* and *x* are plant yield and aboveground biomass, respectively. This method enables geometrical interpretations that lead to statistical testing procedures to compare slope, offset and shift along the allometric line (Warton et al., 2012; Taskinen and Warton, 2013). More specifically, we assessed the effect of plant family and, within each family, the effect of crop type and N fertilization, on the position of individual plants along the main reproductive allometric line. Because of the unequal sample sizes, allometric lines for legumes were compared only i) under non-fertilized conditions, to compare the effect of crop type, and ii) in intercrops, to compare the effect of N fertilization.

When two groups had significantly different slopes of allometric lines, we determined the aboveground biomass for which the two allometric lines intersect, thus defining the plant-size threshold above which a plant had a proportionally higher yield. For example, this threshold equaled the abscissa *X*0 of the intersection of the allometric relationships for an intercrop (IC) and sole crop (SC), calculated as *X*0 = (*aIC* − *aSC*)/(*bSC* − *bIC*), where *aIC* and *aSC* are the estimated intercept, and *bIC* and *bSC* are the least square estimate of the slope of the allometric relationship for an intercrop and sole crop, respectively. We calculated 95% confidence intervals (CI0.95) of this threshold using the procedure of Filliben and McKinney (1972). Moreover, to test the robustness of the differences in slopes, we performed a bootstrap procedure using 10 thousand bootstrap samples for each case study and identified the percentage of cases where the p-value of the sma test was below 0.05 (i.e. indicating a significant difference between the slopes).

To assess the dominance of the focal species in intercrop, we calculated a distance index based on biomass difference (i.e. fitness distance, Mayfield and Levine, 2010; Cadotte, 2017) between the two intercropped species within each of the 159 experimental units in intercropping situations. We first normalized plant yield and biomass values within species *x* fertilization groups to account for major plant size differences between intercropped species (unity-based normalization, *x*′ = (*x* − *x_m_ax*)/*x*_(_*x_m_ax* − *x_m_in*)). Then, considering an intercrop mixing two species *i* and *j*, the biomass distance index for the focal species *i* was defined as *x*′_*j*_ − *x*′_*i*_, and respectively for species *j*. This index ranges from −1 (i.e. focal species is dominant) to +1 (i.e. focal species is dominated). We used a linear model to analyze change in plant yield as a function of the biomass distance index.

The allometric relationships led to centered residuals as the differences between the observed yield and the predicted one from associated biomass. A natural question arose about the impact of the conditions of each experiment on these results. Considering an ANOVA or a mixed model would be unsatisfying due to the unbalanced sizes of each experimental group in the dataset. Thus, an alternative approach was proposed to tackle such a question. The residual values were plotted separately for each subset of data obtained in the same conditions. Moreover, the p-th quantiles for p=2.5% and p=97.5% were drawn to bounds 95% of residual values to visualize possible outliers (Supplementary Material S2). The results indicated that no extreme value appeared as remarkable. Some variability is revealed but its order of magnitude remains below the dispersal of the residuals. The role of the experimental factors appears then as neglectable with respect to the residual variations of the allometric relationships.

Data were analyzed with R software version 3.6.0 with the packages *dplyr* [data processing; Wickham et al. (2019)], *ggplot2* [visualization; Wickham (2016)] and *knitr* [reporting; Xie (2015)].

## Results

### Reproductive allometry in cereals and legumes

The reproductive allometric relationship between plant yield and biomass was significant and robust (R^2^ = 0.94) across all experimental units, indicating that size is a predominant driver of crop yield. Allometric relationships of legumes and cereals displayed a similar slope close to 1 (1.03 ± 0.02), indicating an overall isometric relationship between plant yield and biomass. However, legumes generally had larger biomass and grain yield than cereals (significant shift along the main relationship, Fig. 2A). Moreover, legumes generally had higher yield than cereals for a given biomass (significant offset along the y-axis). The relationships between reproductive effort and plant biomass were weak for both cereals and legumes (R^2^ = 0.104 and 0.049, respectively), with reproductive effort decreasing slightly as plant biomass increased (p < 0.0001, Fig. 2B). Legumes had slightly but significantly higher reproductive effort (0.48 ± 0.10, ranging from 0.19-0.73) than cereals (0.43 ± 0.08, ranging from 0.22-0.67), although it varied greatly.

**Fig. 2.**
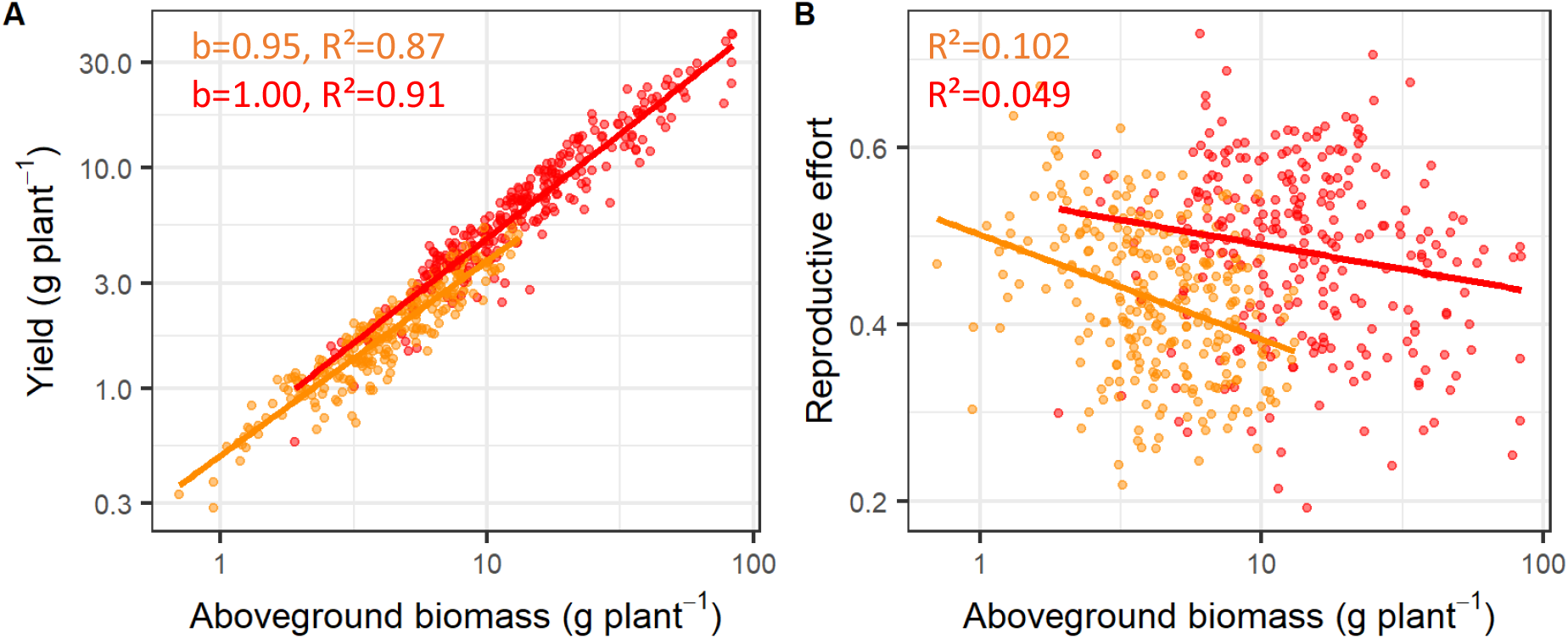
(A) Reproductive allometric relationship (log-log scale) between plant yield (g.plant^−1^) and plant biomass (g.plant^−1^) and (B) relationship between reproductive effort (=yield:biomass) and plant biomass (g.plant^−1^) for cereals (orange) and legumes (red) for all experimental units in intercropping and sole-cropping situations.

### Crop management impacted the plant reproductive allometry, especially for legumes

The slopes of allometric relationships were steeper under intercropping than sole-cropping conditions (Fig. 3A-C). The intercropping effect was stronger in non-fertilized conditions (legumes, p < 10^−6^; then cereals, p = 0.004) than in fertilized conditions (cereals, p = 0.03). However, data points were relatively dispersed but the bootstrap procedure based on 10 thousand bootstrap samples for each modality corroborated these results, i.e. p-value of the *sma* slope test was below 0.05 in 99%, 82% and 61% of the cases among the 10 thousand bootstrap samples, respectively for the legumes in non-fertilized conditions, cereals in non-fertilized conditions and cereals in fertilized conditions. For legumes under non-fertilized conditions, intercrops had significantly higher reproductive effort than sole crops. Although allometric differences were observed for cereals, ANOVAs indicated that intercropping had no significant effect on reproductive effort or its components (plant yield and biomass) whether in fertilized or non-fertilized conditions (Table 2).

**Fig. 3.**
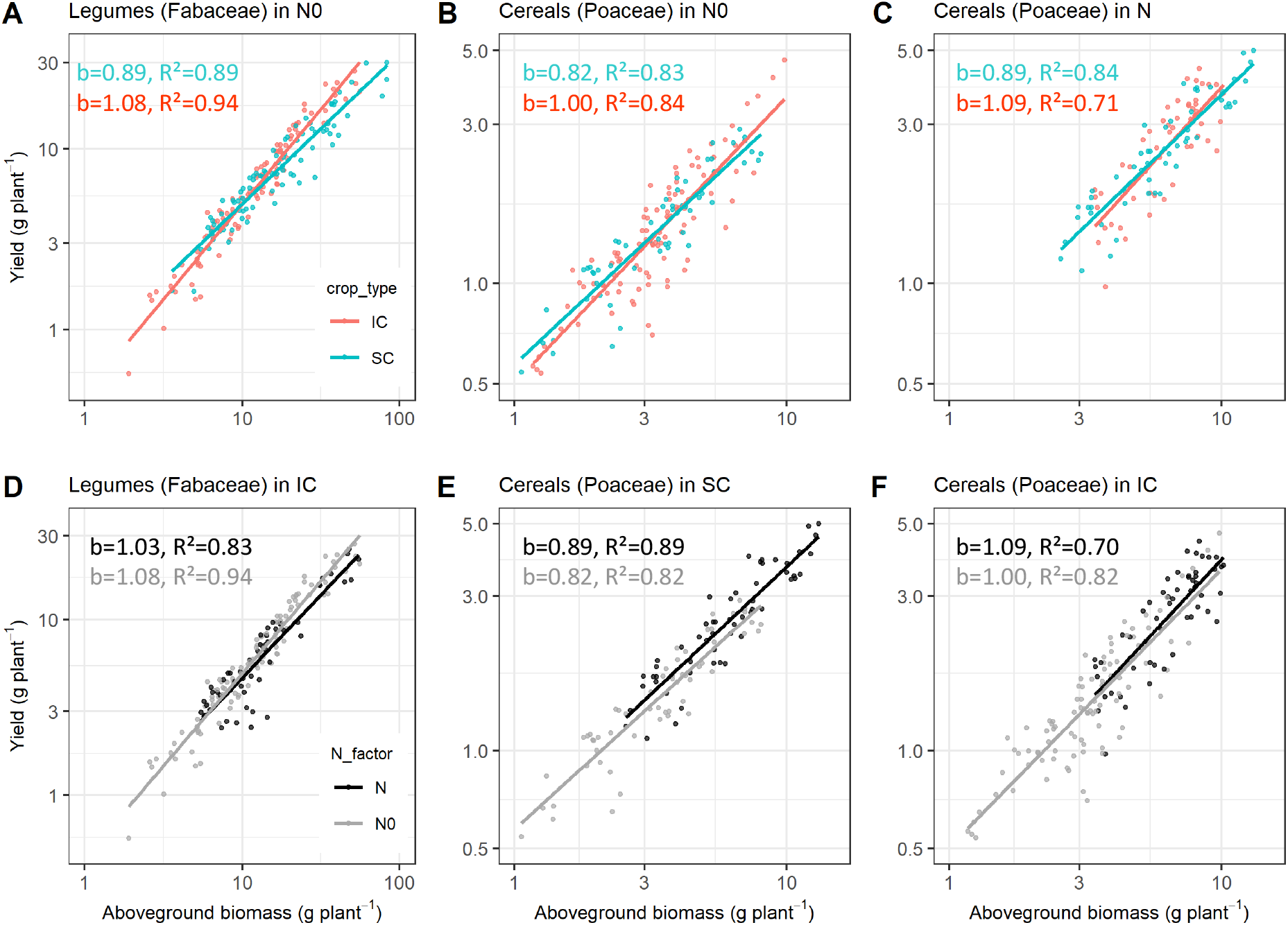
Reproductive allometric relationship (log-log scale) between plant yield (g.plant^−1^) and plant biomass (g.plant^−1^) by i) crop type (i.e. sole crop (SC) *vs.* intercrop (IC)), for (A) legumes (Fabaceae) under non-fertilized conditions (N0), (B) cereals (Poaceae) under N0 and (C) cereals under N-fertilized conditions (N), and by ii) N fertilization, for (D) legumes grown under IC, (E) cereals under SC and (F) cereals under IC. b represents the allometric scaling exponent of the studied relationships.

**Table 2.**
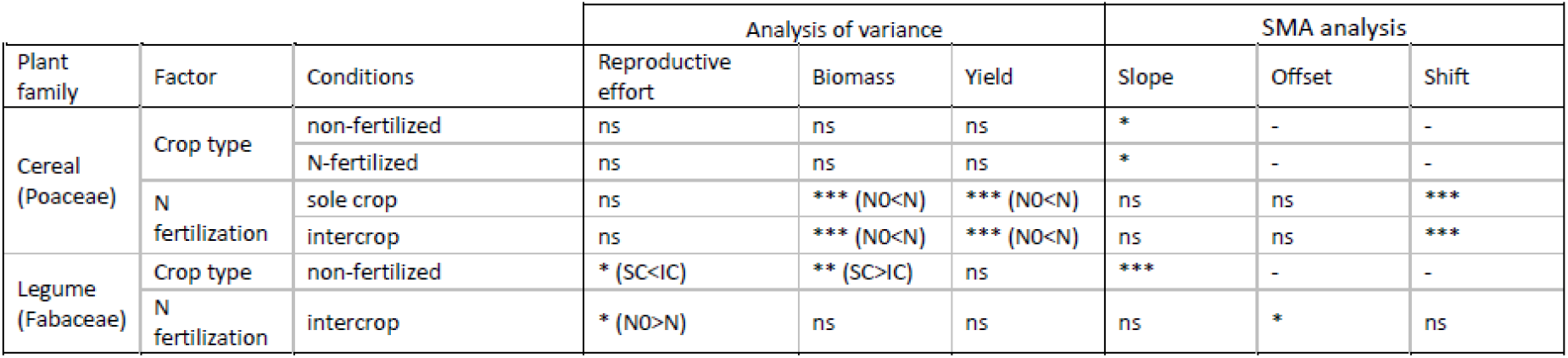
Effect of crop type (sole crop SC *vs.* intercrop IC) and nitrogen (N) fertilization (non-fertilized N0 *vs.* fertilized N) by plant family (Poaceae *vs.* Fabaceae) on i) reproductive effort, plant biomass (g.plant^−1^) and plant yield (g.plant^−1^) using analysis of variance and ii) allometric relationships (slope, offset and shift along the relationship) using standardized major axis (SMA) analysis (*** <0.0001, ** <0.001, * <0.05, ns non-significant).

We calculated the plant size threshold corresponding to the intersection of allometric lines in intercrop and sole crop conditions to identify the minimum plant size corresponding to a positive intercropping effect on biomass allocation (increased yield for a same plant size). The plant size threshold above which a legume under non-fertilized conditions (Fig. 3A) benefited from intercropping was 10.3 g.plant^−1^ (CI0.95 = [6.7-13.5 g.plant^−1^]), with biomass ranging from 1.9-83.2 g.plant^−1^. For a cereal under non-fertilized conditions (Fig. 3B), the threshold was 3.5 g.plant^−1^ (CI0.95 = [2.0-5.9 g.plant^−1^]), with biomass ranging from 0.7-9.9 g.plant^−1^. For a cereal under N-fertilized conditions (Fig. 3C), the threshold was 6.3 g.plant^−1^, with biomass ranging from 2.6-13.2 g.plant^−1^. We could not derive CI0.95 for this situation because the allometric relationships of the two datasets had high collinearity and widely scattered points.

In intercropping conditions, for a given species, we analyzed how variation in its yields depends on biomass distance between the two intercropped species. This distance index significantly explained yield variation in all studied species *x* management cases (R^2^ 0.39 - 0.48, p < 1e-3, Fig. 4).

**Fig. 4.**
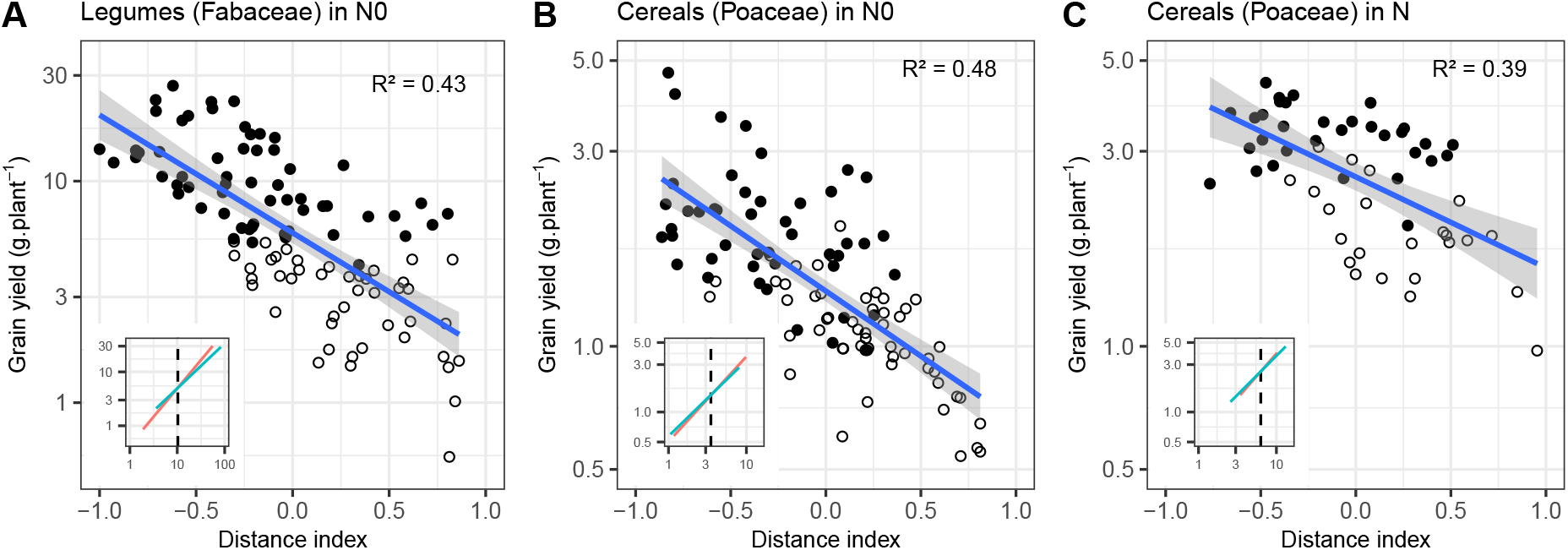
Grain yield (g.plant^−1^) of intercropped components as a function of biomass distance index for (A) legumes (Fabaceae) under non-fertilized conditions (N0), (B) cereals (Poaceae) under N0, and (C) cereals under N-fertilized conditions (N). The distance index was calculated as the biomass difference between the focal species and its associated species. Inset graphs represent reproductive allometric relationships, with vertical dotted lines marking the biomass threshold corresponding to the intersection of allometric lines between sole crops (blue) and intercrops (red). This threshold is used to split the field population where the focal species biomass was above (filled symbols) or below it (open symbols).

N fertilization also influenced allometric relationships (Fig 3D-F). For a given plant biomass, intercropped legumes had higher yield without N fertilization than with it (significant offset, Fig. 3D). The analysis of the reproductive effort confirmed this result, with significantly higher biomass allocation for legumes under non-fertilized (0.51 ± 0.09) than N-fertilized (0.47 ± 0.11) conditions. However, we could not determine whether biomass or yield caused this difference. For cereals, N fertilization did not influence the allometric relationship: N-fertilized plants had proportionally larger biomass and yield, regardless of the crop type (significant shift, Fig. 3E-F). This result was confirmed by both crop types having a similar reproductive effort: both ratio components (plant yield and biomass) were higher under N-fertilized conditions than non-fertilized conditions, regardless of the crop type.

## Discussion

Intercrop design aims to improve resource-use efficiency, especially crop N use (yield per unit of N absorbed) in cereal-legume intercrops (Jensen et al., 2020). In the experiments examined, plant-plant interactions in intercropping conditions influenced reproductive allometry, especially for legumes intercropped with cereals in non-fertilized conditions with an increase of biomass allocation to their reproductive parts. These results increase knowledge about the factors that influence plant allometry while the allometric rules are usually considered invariant across species and ecological situations, notably as expected from the metabolic scaling theory [MST; Niklas and Enquist (2001)]. Moreover MST appears as a new and promising conceptual framework to improve intercrop management. The allometric approach improves our understanding of which intercropping situation favors one species or the other, and provides some guidelines to identify putative trade-offs depending on the agronomic objective assigned to the intercrop (e.g. focus on the legume, or having both intercropped species reaching a suitable yield without one species strongly dominating the other).

In addition to the strong size-dependence of grain yield production, which was predicted by allometric relationships and highlighted in other studies (Sugiyama and Bazzaz, 1998; Vega et al., 2000; Weiner, 2004; Liu et al., 2008; Qin et al., 2013), we observed that species grown in intercrops had a greater increase in yield for a given increase in plant size than species grown in sole crops through the threshold analysis (x-coordinate of the intersection of allometric lines between sole cropping and intercropping conditions). This highlights a better spatial and temporal resource use efficiency in a field situation, which is a pillar of ecological intensification (Li-li et al., 2015).

This effect depends on plant family and the relative dominance of the two intercropped species. When ranking this effect among cropping conditions, intercropping benefited legumes under non-fertilized conditions the most, followed by cereals under non-fertilized conditions and then cereals under N-fertilized conditions. For example, a cereal plant in a sole crop is surrounded by other cereal plants. Since cereal plants generally compete strongly for soil resources, they experience strong intra-specific competition under non-fertilized conditions. If this cereal plant is intercropped with legume plants, however, some of its close neighbors are legumes, which compete less for soil N than cereals (Satorre and Snaydon, 1992; Mariotti et al., 2009) and can fix atmospheric N. This decreases the competition experienced by the cereal plant due to functional complementarity in N acquisition strategy (Hinsinger et al., 2011; Duchene et al., 2017). However, when the intercrop is fertilized with N, cereals have a competitive advantage over legumes and complementarity for resource use is replaced with strong interspecific competition from the cereal over the legume. Thus, intercrop design should focus on two key points: i) plant family, characterized by their competitive ability (Goldberg, 1990); and ii) characteristics of the two intercropped species and their capacity to compete for limiting resources depending on the plant neighborhood (Stoll and Weiner, 2000; Gaudio et al., 2019).

The relaxation of competitive interaction in intercropping situations is highlighted by the strong effect of biomass differences between the two intercropped species in grain yield production [also called fitness distance in the ecological literature; Cadotte (2017)]. This reflects the difference in dominance between the two intercropped species (Mayfield and Levine, 2010), which is one key driver for competitive exclusion. Then, in intercrops, yield of cereals in N-fertilized conditions is hardly influenced by the biomass of the associated legume, whereas yield of cereals and legumes in non-fertilized conditions were much more influenced by the biomass of the associated species. When the biomass difference between the two intercropped species is high, there is an obvious imbalance between the two species, leading to strong differences in competitive ability of the two components of the mixture: the greater the biomass difference, the more intense hierarchical competition (Kunstler et al., 2016). Therefore, around the size threshold corresponding to the intersection between allometric lines in sole- and intercropping conditions identified for each cropping situation, the two intercropped species do not reach their maximal size and associated yield but an equilibrium exists between them. When we move away from this threshold, one intercrop component becomes highly dominant or conversely dominated due to hierarchical competition.

For legumes, we showed that fertilization decreases the reproductive effort (i.e. lower yield for a given biomass) compared to that under non-fertilized intercropping situations, as highlighted in other studies (e.g. Corre-Hellou et al., 2007 for spring barley-pea). This is probably because when strong competition limits a resource, plants tend to allocate more biomass to structures associated with acquiring it, potentially to the detriment of reproductive organs (Poorter and Nagel, 2000; Bonser, 2013). In contrast, cereals in intercrops had similar reproductive effort whatever the fertilization condition, but a given reproductive effort was reached with proportionally higher biomass and yield in fertilized plots. In these cropping conditions, cereals are larger and compete more for aboveground and soil resources. Understanding the causal processes for such a modified allocation would require targeted experiments that measure key functional traits of legumes that reflect plant behavior for light (e.g. specific leaf area) and N availability (e.g. leaf N content) (Freschet et al., 2018).

## Pathway to applications

Weiner et al. (2009) described the reproductive-vegetative allometric relationship as a relatively fixed-boundary condition, meaning that a plant cannot increase its reproductive output without growing more first. Our findings highlight that, in a crop mixture, the interaction between the two intercropped species makes this boundary more complex given that the reproductive output of one component also depends on the performance of the other. From a practical viewpoint, the analysis of the intersection of allometric relationships enables the identification of plant biomass thresholds for each component of the mixture. We propose to use these thresholds as management criteria to cultivate each species up to its reproductive output maximum while accounting for the performance of the other.

We found that plant-plant interactions were a strong driver of yield variation in crop mixtures. In this case, an agronomic action on one species can readily influence the other even under relatively variable climate conditions, which is an important observation for designing and managing intercrops. For instance, if the goal of the farmer is to promote legume yield, using the cereal mainly to limit legume disease and lodging (e.g. Viguier et al., 2018 for spring wheat-lentil intercrops), then we should identify plant size level of the target legume above which higher growth means higher yield, accounting for the interaction with the cereal. Conversely, if the goal is to promote both intercropped species (Hauggaard-Nielsen et al., 2008; Pelzer et al., 2014), then trade-offs should be managed to be close to the threshold in order to avoid a strong dominance of one of the two species. However, while such ecological knowledge is important for informing a sustainable transition, we argue that many other factors influence decision-making beyond scientific evidence alone (Hazard et al., 2020).

Our results indicated that legumes would better benefit from intercropping, which is aligned with the need to increase legumes acreage, a stake with both environmental and food aspects. On one hand, greenhouse gas emissions can be reduced through a decreased use of synthetic N fertilizer (Zander et al., 2016) and a shift to plant-based protein diets. On the other hand, increasing plant-based protein production to sustain a nutritional transition is limited by legumes productivity in sole crop (Magrini et al., 2018). In any case, intercropping appears as an additional lever in the agro-ecological transition.

## Authors’ contributions

NG, PC, CV, FF designed and planned the study;

LB, GCH, AC, HHN, PH, ESJ, EPJ, EJ, BK, CN, EP provided data;

NG, PC, CB, RM designed, formatted and homogenized the database;

NG, PC, XG, RM analyzed the data;

NG, PC, CV, FF, EP, SM, XG, RM, HHN, LB, AC, CN, NM participated in the writing and editing.

## Acknowledgements

The authors thank all the technical staff who helped to acquire this vast dataset, Michael and Michelle Corson for their helpful comments and English revision, and the reviewers and editor for their contribution to highly improve this manuscript. This study was supported by the French National Research Agency under the Investments for the Future Program (ANR-16-CONV-0004) and by the INRA Environment and Agronomy Division through the project IDEA (Intra- and interspecific diversity mixture in agriculture). CV was supported by the European Research Council (ERC) Starting Grant Project ‘ecophysiological and biophysical constraints on domestication in crop plants’ (grant ERC-StG-2014-639706-CONSTRAINTS).

## Data availability statement

The data that support the findings of this study are openly available at Dryad Digital Repository: https://doi.org/10.5061/dryad.9ghx3ffhv (Gaudio et al., 2021).

## References

Bedoussac, L., and E. Justes. 2010b. Dynamic analysis of competition and complementarity for light and N use to understand the yield and the protein content of a durum wheat–winter pea intercrop. Plant and Soil 330(1–2): 37–54. doi: 10.1007/s11104-010-0303-8.

Bedoussac, L., and E. Justes. 2010a. The efficiency of a durum wheat-winter pea intercrop to improve yield and wheat grain protein concentration depends on N availability during early growth. Plant and Soil 330(1–2): 19–35. doi: 10.1007/s11104-009-0082-2.

Beillouin, D., T. Ben-Ari, and D. Makowski. 2019. A dataset of meta-analyses on crop diversification at the global scale. Data in Brief 24: 103898. doi: 10.1016/j.dib.2019.103898.

Bonser, S.P. 2013. High reproductive efficiency as an adaptive strategy in competitive environments. Functional Ecology 27(4): 876–885. doi: 10.1111/1365-2435.12064.

Cadotte, M.W. 2017. Functional traits explain ecosystem function through opposing mechanisms. Ecology Letters 20(8): 989–996. doi: 10.1111/ele.12796.

Cheplick, G.P. 2005. The Allometry of Reproductive Allocation. In: Reekie, E.G. and Bazzaz, F.A., editors, Reproductive Allocation in Plants. Academic Press, Burlington. p. 97–128

Coomes, D.A., E.R. Lines, and R.B. Allen. 2011. Moving on from Metabolic Scaling Theory: Hierarchical models of tree growth and asymmetric competition for light. Journal of Ecology 99(3): 748–756. doi: 10.1111/j.1365-2745.2011.01811.x.

Corre-Hellou, G., N. Brisson, M. Launay, J. Fustec, and Y. Crozat. 2007. Effect of root depth penetration on soil nitrogen competitive interactions and dry matter production in pea-barley intercrops given different soil nitrogen supplies. Field Crops Research 103(1): 76–85. doi: 10.1016/j.fcr.2007.04.008.

Corre-Hellou, G., J. Fustec, and Y. Crozat. 2006. Interspecific competition for soil N and its interaction with N-2 fixation, leaf expansion and crop growth in pea-barley intercrops. Plant and Soil 282(1–2): 195–208. doi: 10.1007/s11104-005-5777-4.

Deng, J., J. Ran, Z. Wang, Z. Fan, G. Wang, et al. 2012. Models and tests of optimal density and maximal yield for crop plants. Proceedings of the National Academy of Sciences of the United States of America 109(39): 15823–15828. doi: 10.1073/pnas.1210955109.

Duchene, O., J.-F. Vian, and F. Celette. 2017. Intercropping with legume for agroecological cropping systems: Complementarity and facilitation processes and the importance of soil microorganisms. A review. Agriculture, Ecosystems & Environment 240(Supplement C): 148–161. doi: 10.1016/j.agee.2017.02.019.

Echarte, L., and F.H. Andrade. 2003. Harvest index stability of Argentinean maize hybrids released between 1965 and 1993. Field Crops Research 82(1): 1–12. doi: 10.1016/S0378-4290(02)00232-0.

Enquist, B.J., G.B. West, E.L. Charnov, and J.H. Brown. 1999. Allometric scaling of production and life-history variation in vascular plants. Nature 401(6756): 907–911. doi: 10.1038/44819.

Filliben, J.J., and J.E. McKinney. 1972. Confidence limits for the abscissa of intersection of two linear regressions. Journal of Research of the National Bureau of Standards -B. Mathematical Sciences 3-4(76B): 179–192. http://archive.org/details/jresv76Bn3-4p179 (accessed 15 October 2020).

Freschet, G.T., C. Violle, M.Y. Bourget, M. Scherer-Lorenzen, and F. Fort. 2018. Allocation, morphology, physiology, architecture: The multiple facets of plant above- and below-ground responses to resource stress. New Phytologist 219(4): 1338–1352. doi: 10.1111/nph.15225.

Gaba, S., A. Alignier, S. Aviron, S. Barot, M. Blouin, et al. 2018. Ecology for Sustainable and Multifunctional Agriculture. In: Gaba, S., Smith, B., and Lichtfouse, E., editors, Sustainable Agriculture Reviews 28: Ecology for Agriculture. p. 1–46

Gaudio, N., A.J. Escobar-Gutiérrez, P. Casadebaig, J.B. Evers, F. Gérard, et al. 2019. Current knowledge and future research opportunities for modeling annual crop mixtures. A review. Agronomy for Sustainable Development 39(2): 20. doi: 10.1007/s13593-019-0562-6.

Gaudio, N., C. Violle, X. Gendre, F. Fort, R. Mahmoud, et al. 2021. Interspecific interactions regulate plant reproductive allometry in cereal-legume intercropping systems. doi: 10.5061/DRYAD.9GHX3FFHV.

Goldberg, D.E. 1990. Components of resource competition in plant communities. In: Grace, J.B. and Tilman, D., editors, Perspectives on plant competition. p. 27–49

Hauggaard-Nielsen, H., M. Gooding, P. Ambus, G. Corre-Hellou, Y. Crozat, et al. 2009a. Pea-barley intercropping for efficient symbiotic N-2-fixation, soil N acquisition and use of other nutrients in European organic cropping systems. Field Crops Research 113(1): 64–71. doi: 10.1016/j.fcr.2009.04.009.

Hauggaard-Nielsen, H., M. Gooding, P. Ambus, G. Corre-Hellou, Y. Crozat, et al. 2009b. Pea-barley intercropping and short-term subsequent crop effects across European organic cropping conditions. Nutrient Cycling in Agroecosystems 85(2): 141–155. doi: 10.1007/s10705-009-9254-y.

Hauggaard-Nielsen, H., B. Jørnsgaard, J. Kinane, and E.S. Jensen. 2008. Grain legume–cereal intercropping: The practical application of diversity, competition and facilitation in arable and organic cropping systems. Renewable Agriculture and Food Systems 23(1): 3–12. doi: 10.1017/S1742170507002025.

Hay, R. 1995. Harvest Index - a Review of Its Use in Plant-Breeding and Crop Physiology. Annals of Applied Biology 126(1): 197–216. doi: 10.1111/j.1744-7348.1995.tb05015.x.

Hazard, L., M. Cerf, C. Lamine, D. Magda, and P. Steyaert. 2020. A tool for reflecting on research stances to support sustainability transitions. Nature Sustainability 3(2): 89–95. doi: 10.1038/s41893-019-0440-x.

Hinsinger, P., E. Betencourt, L. Bernard, A. Brauman, C. Plassard, et al. 2011. P for Two, Sharing a Scarce Resource: Soil Phosphorus Acquisition in the Rhizosphere of Intercropped Species. Plant Physiology 156(3): 1078–1086. doi: 10.1104/pp.111.175331.

Jensen, E.S., G. Carlsson, and H. Hauggaard-Nielsen. 2020. Intercropping of grain legumes and cereals improves the use of soil N resources and reduces the requirement for synthetic fertilizer N: A global-scale analysis. Agronomy for Sustainable Development 40(1): 5. doi: 10.1007/s13593-020-0607-x.

Knudsen, M.T., H. Hauggaard-Nielsen, B. Jornsgard, and E.S. Jensen. 2004. Comparison of interspecific competition and N use in pea-barley, faba bean-barley and lupin-barley intercrops grown at two temperate locations. Journal of Agricultural Science 142: 617–627. doi: 10.1017/S0021859604004745.

Kunstler, G., D. Falster, D.A. Coomes, F. Hui, R.M. Kooyman, et al. 2016. Plant functional traits have globally consistent effects on competition. Nature 529(7585): 204–U174. doi: 10.1038/nature16476.

Launay, M., N. Brisson, S. Satger, H. Hauggaard-Nielsen, G. Corre-Hellou, et al. 2009. Exploring options for managing strategies for pea-barley intercropping using a modeling approach. European Journal of Agronomy 31(2): 85–98. doi: 10.1016/j.eja.2009.04.002.

Li-li, M., Z. Li-zhen, Z. Si-ping, J.B. Evers, W. van der Werf, et al. 2015. Resource use efficiency, ecological intensification and sustainability of intercropping systems. Journal of Integrative Agriculture 14(8): 1542–1550. doi: 10.1016/S2095-3119(15)61039-5.

Lin, B.B. 2011. Resilience in Agriculture through Crop Diversification: Adaptive Management for Environmental Change. Bioscience 61(3): 183–193. doi: 10.1525/bio.2011.61.3.4.

Litrico, I., and C. Violle. 2015. Diversity in Plant Breeding A New Conceptual Framework. Trends in Plant Science 20(10): 604–613. doi: 10.1016/j.tplants.2015.07.007.

Liu, J., G.-X. Wang, L. Wei, and C.-M. Wang. 2008. Reproductive allocation patterns in different density populations of spring wheat. Journal of Integrative Plant Biology 50(2): 141–146. doi: 10.1111/j.1744-7909.2007.00602.x.

Magrini, M.-B., M. Anton, J.-M. Chardigny, G. Duc, M. Duru, et al. 2018. Pulses for Sustainability: Breaking Agriculture and Food Sectors Out of Lock-In. Frontiers in Sustainable Food Systems 2: UNSP 64. doi: 10.3389/fsufs.2018.00064.

Malézieux, E. 2012. Designing cropping systems from nature. Agronomy for Sustainable Development 32(1): 15–29. doi: 10.1007/s13593-011-0027-z.

Malézieux, E., Y. Crozat, C. Dupraz, M. Laurans, D. Makowski, et al. 2009. Mixing plant species in cropping systems: Concepts, tools and models. A review. Agronomy for Sustainable Development 29(1): 43–62. doi: 10.1051/agro:2007057.

Mariotti, M., A. Masoni, L. Ercoli, and I. Arduini. 2009. Above- and below-ground competition between barley, wheat, lupin and vetch in a cereal and legume intercropping system. Grass and Forage Science 64(4): 401–412. doi: 10.1111/j.1365-2494.2009.00705.x.

Mayfield, M.M., and J.M. Levine. 2010. Opposing effects of competitive exclusion on the phylogenetic structure of communities. Ecology Letters 13(9): 1085–1093. doi: 10.1111/j.1461-0248.2010.01509.x.

Milla, R., C.P. Osborne, M.M. Turcotte, and C. Violle. 2015. Plant domestication through an ecological lens. Trends in Ecology & Evolution 30(8): 463–469. doi: 10.1016/j.tree.2015.06.006.

Naudin, C., G. Corre-Hellou, S. Pineau, Y. Crozat, and M.-H. Jeuffroy. 2010. The effect of various dynamics of N availability on winter pea-wheat intercrops: Crop growth, N partitioning and symbiotic N-2 fixation. Field Crops Research 119(1): 2–11. doi: 10.1016/j.fcr.2010.06.002.

Naudin, C., H.M.G. van der Werf, M.-H. Jeuffroy, and G. Corre-Hellou. 2014. Life cycle assessment applied to pea-wheat intercrops: A new method for handling the impacts of co-products. Journal of Cleaner Production 73: 80–87. doi: 10.1016/j.jclepro.2013.12.029.

Nee, S., N. Colegrave, S.A. West, and A. Grafen. 2005. The illusion of invariant quantities in life histories. Science 309(5738): 1236–1239. doi: 10.1126/science.1114488.

Niklas, K.J., and B.J. Enquist. 2001. Invariant scaling relationships for interspecific plant biomass production rates and body size. Proceedings of the National Academy of Sciences of the United States of America 98(5): 2922–2927. doi: 10.1073/pnas.041590298.

Pelzer, E., M. Bazot, L. Guichard, and M.-H. Jeuffroy. 2016. Crop Management Affects the Performance of a Winter Pea-Wheat Intercrop. Agronomy Journal 108(3): 1089–1100. doi: 10.2134/agronj2015.0440.

Pelzer, E., M. Bazot, D. Makowski, G. Corre-Hellou, C. Naudin, et al. 2012. Pea–wheat intercrops in low-input conditions combine high economic performances and low environmental impacts. European Journal of Agronomy 40(Supplement C): 39–53. doi: 10.1016/j.eja.2012.01.010.

Pelzer, E., N. Hombert, M.-H. Jeuffroy, and D. Makowski. 2014. Meta-Analysis of the Effect of Nitrogen Fertilization on Annual Cereal-Legume Intercrop Production. Agronomy Journal 106(5): 1775–1786. doi: 10.2134/agronj13.0590.

Poorter, H., A.M. Jagodzinski, R. Ruiz-Peinado, S. Kuyah, Y. Luo, et al. 2015. How does biomass distribution change with size and differ among species? An analysis for 1200 plant species from five continents. New Phytologist 208(3): 736–749. doi: 10.1111/nph.13571.

Poorter, H., and O. Nagel. 2000. The role of biomass allocation in the growth response of plants to different levels of light, CO2, nutrients and water: A quantitative review. Australian Journal of Plant Physiology 27(6): 595–607. doi: 10.1071/PP99173.

Poorter, H., and L. Sack. 2012. Pitfalls and possibilities in the analysis of biomass allocation patterns in plants. Frontiers in Plant Science 3: 259. doi: 10.3389/fpls.2012.00259.

Qin, X., J. Weiner, L. Qi, Y. Xiong, and F. Li. 2013. Allometric analysis of the effects of density on reproductive allocation and Harvest Index in 6 varieties of wheat (Triticum). Field Crops Research 144: 162–166. doi: 10.1016/j.fcr.2012.12.011.

R Core Team. 2019. R: A Language and Environment for Statistical Computing. R Foundation for Statistical Computing, Vienna, Austria.

Satorre, E.H., and R.W. Snaydon. 1992. A comparison of root and shoot competition between spring cereals and Avena fatua L. Weed Research 32(1): 45–55. doi: 10.1111/j.1365-3180.1992.tb01861.x.

Stoll, P., and J. Weiner. 2000. A neighborhood view of interactions among individual plants. In: Dieckmann, U., Law, R., and Metz, J.A.J., editors, The geometry of ecological interactions: Simplifying spatial complexity. Cambridge University Press. p. 11–27

Sugiyama, S., and F.A. Bazzaz. 1998. Size dependence of reproductive allocation: The influence of resource availability, competition and genetic identity. Functional Ecology 12(2): 280–288. doi: 10.1046/j.1365-2435.1998.00187.x.

Tang, X., S.A. Placella, F. Dayde, L. Bernard, A. Robin, et al. 2016. Phosphorus availability and microbial community in the rhizosphere of intercropped cereal and legume along a P-fertilizer gradient. Plant and Soil 407(1–2): 119–134. doi: 10.1007/s11104-016-2949-3.

Taskinen, S., and D.I. Warton. 2013. Robust tests for one or more allometric lines. Journal of Theoretical Biology 333: 38–46. doi: 10.1016/j.jtbi.2013.05.010.

Thorsted, M.D., J. Weiner, and J.E. Olesen. 2006. Above- and below-ground competition between intercropped winter wheat Triticum aestivum and white clover Trifolium repens. Journal of Applied Ecology 43(2): 237–245. doi: 10.1111/j.1365-2664.2006.01131.x.

Unkovich, M., J. Baldock, and M. Forbes. 2010. Advances in Agronomy. In: Sparks, D.L., editor. p. 173–219

Vasseur, F., M. Exposito-Alonso, O.J. Ayala-Garay, G. Wang, B.J. Enquist, et al. 2018. Adaptive diversification of growth allometry in the plant Arabidopsis thaliana. Proceedings of the National Academy of Sciences 115(13): 3416–3421. doi: 10.1073/pnas.1709141115.

Vasseur, F., C. Violle, B.J. Enquist, C. Granier, and D. Vile. 2012. A common genetic basis to the origin of the leaf economics spectrum and metabolic scaling allometry. Ecology Letters 15(10): 1149–1157. doi: 10.1111/j.1461-0248.2012.01839.x.

Vega, C.R.C., V.O. Sadras, F.H. Andrade, and S.A. Uhart. 2000. Reproductive allometry in soybean, maize and sunflower. Annals of Botany 85(4): 461–468. doi: 10.1006/anbo.1999.1084.

Viguier, L., L. Bedoussac, E.-P. Journet, and E. Justes. 2018. Yield gap analysis extended to marketable grain reveals the profitability of organic lentil-spring wheat intercrops. Agronomy for Sustainable Development 38(4): 39. doi: 10.1007/s13593-018-0515-5.

Warton, D.I., R.A. Duursma, D.S. Falster, and S. Taskinen. 2012. Smatr 3-an R package for estimation and inference about allometric lines. Methods in Ecology and Evolution 3(2): 257–259. doi: 10.1111/j.2041-210X.2011.00153.x.

Warton, D.I., I.J. Wright, D.S. Falster, and M. Westoby. 2006. Bivariate line-fitting methods for allometry. Biological Reviews 81(2): 259–291. doi: 10.1017/S1464793106007007.

Weiner, J. 2004. Allocation, plasticity and allometry in plants. Perspectives in Plant Ecology Evolution and Systematics 6(4): 207–215. doi: 10.1078/1433-8319-00083.

Weiner, J., L.G. Campbell, J. Pino, and L. Echarte. 2009. The allometry of reproduction within plant populations. Journal of Ecology 97(6): 1220–1233. doi: 10.1111/j.1365-2745.2009.01559.x.

West, G.B., J.H. Brown, and B.J. Enquist. 1997. A general model for the origin of allometric scaling laws in biology. Science 276(5309): 122–126. doi: 10.1126/science.276.5309.122.

West, G.B., J.H. Brown, and B.J. Enquist. 1999. A general model for the structure and allometry of plant vascular systems. Nature 400(6745): 664–667.

Wickham, H. 2016. ggplot2: Elegant Graphics for Data Analysis. 2nd ed. Springer International Publishing.

Wickham, H., R. François, L. Henry, and K. Müller. 2019. Dplyr: A Grammar of Data Manipulation.

Willey, R.W. 1979. Intercropping: Its importance and research needs. Part 1, competition and yield advantages. Commonwealth Agricultural Bureaux, Place of publication not identified.

Xie, Y. 2015. Dynamic Documents with R and knitr, Second Edition. 2nd ed. Routledge, Boca Raton.

Yu, Y., T.-J. Stomph, D. Makowski, L. Zhang, and W. van der Werf. 2016. A meta-analysis of relative crop yields in cereal/legume mixtures suggests options for management. Field Crops Research 198: 269–279. doi: 10.1016/j.fcr.2016.08.001.

Zander, P., T.S. Amjath-Babu, S. Preissel, M. Reckling, A. Bues, et al. 2016. Grain legume decline and potential recovery in European agriculture: A review. Agronomy for Sustainable Development 36(2): 26. doi: 10.1007/s13593-016-0365-y.

