## supplementary table 1 for "Interspecific interactions regulate plant reproductive allometry in cereal-legume intercropping systems"

Appendix S1. Description of the data used. SC = sole crop; IC = intercrop; N = nitrogen

| Intercropped species (cereal – legume) | Site | Year | No. genotypes (cereal - legume) | IC spatial pattern | Intercrop density (% of the sole crop) | N-treatments (in addition to N0 for all experiments and modalities) | References |
|---|---|---|---|---|---|---|---|
| Spring barley – faba bean | Denmark, Copenhagen | 2001 | 2 - 1 | within-row | 0.5 - 0.5 | 1 N-treatment for barley in SC (50 kg N.ha$^{-1}$) | (Knudsen et al., 2004; Hauggaard-Nielsen et al., 2008) |
| | Denmark, Jyndevad | 2001, 2002, 2003 | | | | | |
| Spring barley - pea | Denmark, Copenhagen | 2001 | 2 - 2 | within-row | 0.5 - 0.5 | 1 N-treatment for barley in SC (50 kg N.ha$^{-1}$) | (Knudsen et al., 2004; Hauggaard-Nielsen et al., 2008) |
| | Denmark, Jyndevad | 2001, 2002, 2003 | | | | | |
| | France, Angers (FNAMS) | 2003 | 1 - 1 | alternate rows | 0.5 - 0.5 | 1 N-treatment (130 kg N.ha$^{-1}$) | (Corre-Hellou et al., 2006) |
| | Denmark, Taastrup | 2003 | 1 - 1 | alternate rows | 0.5 - 0.5 | | (Hauggaard-Nielsen et al., 2009a; b; Launay et al., 2009) |
| | France, Angers (Thorigné) | 2003, 2004 | | | | | |
| | Germany, Kassel | 2004 | | | | | |
| | Italy, San Marco Argentano | 2003, 2004 | | | | | |
| | United Kingdom, Reading | 2003 | | | | | |
| Winter durum wheat – faba bean | France, Auzeville | 2010 | 1 - 1 | alternate rows; within-row | 0.33 - 0.5; 0.5 - 0.5; 0.67 - 0.5 | 3 N-treatments (60, 80, 140 kg N.ha$^{-1}$) | |
| | France, Auzeville (PK) | 2011 | 1 - 1 | alternate rows | 0.5 - 0.5 | (3 P-treatments: P0, 11 and 33 kg P.ha$^{-1}$) | (Tang et al., 2016) |
| | France, Auzeville (PP) | 2011 | 1 - 1 | within-row | 0.5 - 0.5 | 1 N-treatment (140 kg N.ha$^{-1}$) | |
| | France, Auzeville | 2012 | 3 - 4 | within-row | 0.5 - 0.5 | | (Kammoun, 2014) |
| | France, Auzeville | 2013 | 3 - 4 | within-row | 0.5 - 0.5 | 1 N-treatment for wheat in SC (140 kg N.ha$^{-1}$) | (Kammoun, 2014) |
| Winter durum wheat - pea | France, Auzeville | 2006 | 1 - 1 | alternate rows | 0.5 - 0.5 | 2 N-treatments (100, 180 kg N.ha$^{-1}$), except for pea in SC | (Bedoussac and Justes, 2010a; b) |
| | France, Auzeville | 2007 | 4 - 1 | alternate rows | 0.5 - 0.5 | 3 N-treatments (60, 80, 140 kg N.ha$^{-1}$), except for pea in SC | (Bedoussac and Justes, 2010a; b) |
| | France, Auzeville | 2012 | 3 - 4 | within-row | 0.5 - 0.5 | | (Kammoun, 2014) |
| | France, Auzeville | 2013 | 3 - 5 | within-row | 0.5 - 0.5 | 1 N-treatment (140 kg N.ha$^{-1}$), except for pea in SC | (Kammoun, 2014) |
| | France, Auzeville | 2015 | 1 - 4 | within-row | 0.5 - 0.5 | | |
| Winter soft wheat - pea | France, Angers (Thorigné) | 2006 | 1 - 1 | within-row | 0.3 - 0.7; 0.5 - 0.5 | | |
| | France, Angers (Thorigné) | 2007 | 1 - 1 | within-row | 0.5 - 0.5; 0.7 - 0.3 | 1 N-treatment (30 kg N.ha$^{-1}$), except for pea in SC | |
| | France, Angers (La Jaillière) | 2007, 2008 | 1 - 1 | within-row | 0.5 - 0.5 | 4 N-treatments:<br>- 45 kg N.ha$^{-1}$ for all IC and wheat in SC<br>- 190 kg N.ha$^{-1}$ for wheat in SC<br>- 45 (later in the cycle) and 30 kg N.ha$^{-1}$ for all IC | (Naudin et al., 2010, 2014) |
| | France, Angers (Thorigné) | 2008 | 1 - 1 | within-row | 0.5 - 0.5; 0.7 - 0.3 | 3 N-treatments:<br>- 110 kg N.ha$^{-1}$ for wheat in SC<br>- 35 and 72 kg N.ha$^{-1}$ for all IC | |
| | France, Angers (Thorigné) | 2009 | 1 - 1 | within-row | 0.5 - 0.5; 0.7 - 0.3 | 1 N-treatment (40 g N.ha-1), except for pea SC | |
| | France, Grignon | 2010 | 1 - 1 | within-row | 0.33 - 0.66; 0.5 - 0.5; 0.7 - 0.5 | 5 N-treatments:<br>- 200 kg N.ha$^{-1}$ for wheat in SC<br>- 90 kg N.ha$^{-1}$ for wheat in SC and all IC<br>- 140 kg N.ha$^{-1}$ for IC 0.5-0.5 (end of winter or flowering)<br>- 45 kg N.ha$^{-1}$ for IC 0.33-0.66 | (Pelzer et al., 2016) |
| | France, Grignon | 2017 | 1 - 2 | within-row | 0.5 - 0.5 | | |

### References


Bedoussac, L., & Justes, E. (2010a). Dynamic analysis of competition and complementarity for light and N use to understand the yield and the protein content of a durum wheat–winter pea intercrop. *Plant and Soil*, *330*(1–2), 37–54. doi: 10.1007/s11104-010-0303-8

Bedoussac, L., & Justes, E. (2010b). The efficiency of a durum wheat-winter pea intercrop to improve yield and wheat grain protein concentration depends on N availability during early growth. *Plant and Soil*, *330*(1–2), 19–35. doi: 10.1007/s11104-009-0082-2

Corre-Hellou, G., Fustec, J., & Crozat, Y. (2006). Interspecific competition for soil N and its interaction with N-2 fixation, leaf expansion and crop growth in pea-barley intercrops. *Plant and Soil*, *282*(1–2), 195–208. doi: 10.1007/s11104-005-5777-4

Hauggaard-Nielsen, H., Gooding, M., Ambus, P., Corre-Hellou, G., Crozat, Y., Dahlmann, C., … Jensen, E. S. (2009). Pea-barley intercropping and short-term subsequent crop effects across European organic cropping conditions. *Nutrient Cycling in Agroecosystems*, *85*(2), 141–155. doi: 10.1007/s10705-009-9254-y

Hauggaard-Nielsen, H., Gooding, M., Ambus, P., Corre-Hellou, G., Crozat, Y., Dahlmann, C., … Jensen, E. S. (2009). Pea-barley intercropping for efficient symbiotic N-2-fixation, soil N acquisition and use of other nutrients in European organic cropping systems. *Field Crops Research*, *113*(1), 64–71. doi: 10.1016/j.fcr.2009.04.009

Hauggaard-Nielsen, H., Jørnsgaard, B., Kinane, J., & Jensen, E. S. (2008). Grain legume–cereal intercropping: The practical application of diversity, competition and facilitation in arable and organic cropping systems. *Renewable Agriculture and Food Systems*, *23*(1), 3–12. doi: 10.1017/S1742170507002025

Kammoun, B. (2014). *Analyse des interactions génotype x environnement x conduite culturale de peuplement bi-spécifique de cultures associées de blé dur et de légumineuses à graines, à des fins de choix variétal et d'optimisation de leurs itinéraires techniques* (PhD Thesis, Toulouse, INPT). Toulouse, INPT, Toulouse, France. Retrieved from http://www.theses.fr/2014INPT0139

Knudsen, M. T., Hauggaard-Nielsen, H., Jornsgard, B., & Jensen, E. S. (2004). Comparison of interspecific competition and N use in pea-barley, faba bean-barley and lupin-barley intercrops grown at two temperate locations. *Journal of Agricultural Science*, *142*, 617–627. doi: 10.1017/S0021859604004745

Launay, M., Brisson, N., Satger, S., Hauggaard-Nielsen, H., Corre-Hellou, G., Kasynova, E., … Gooding, M. J. (2009). Exploring options for managing strategies for pea-barley intercropping using a modeling approach. *European Journal of Agronomy*, *31*(2), 85–98. doi: 10.1016/j.eja.2009.04.002

Naudin, C., Corre-Hellou, G., Pineau, S., Crozat, Y., & Jeuffroy, M.-H. (2010). The effect of various dynamics of N availability on winter pea-wheat intercrops: Crop growth, N partitioning and symbiotic N-2 fixation. *Field Crops Research*, *119*(1), 2–11. doi: 10.1016/j.fcr.2010.06.002

Pelzer, E., Bazot, M., Guichard, L., & Jeuffroy, M.-H. (2016). Crop Management Affects the Performance of a Winter Pea-Wheat Intercrop. *Agronomy Journal*, *108*(3), 1089–1100. doi: 10.2134/agronj2015.0440

Tang, X., Placella, S. A., Dayde, F., Bernard, L., Robin, A., Journet, E.-P., … Hinsinger, P. (2016). Phosphorus availability and microbial community in the rhizosphere of intercropped cereal and legume along a P-fertilizer gradient. *Plant and Soil*, *407*(1–2), 119–134. doi: 10.1007/s11104-016-2949-3
