## supplementary figure 1 for "Interspecific interactions regulate plant reproductive allometry in cereal-legume intercropping systems"

Appendix S2. Residual distribution of allometric relationships depending on environment (site x year).

The residual values of allometric relationships were retrieved and plotted separately for each subset of data obtained in the same conditions (site). The p-th quantiles for p=2.5% and p=97.5% were drawn (red dotted lines) to bounds 95% of residual values to visualize possible outliers.
The results indicated that no extreme value appeared as remarkable. Some variability is revealed but its order of magnitude remains below the dispersal of the residuals. The role of the experimental factors appears then as neglectable with respect to the residual variations of the linear relationships.

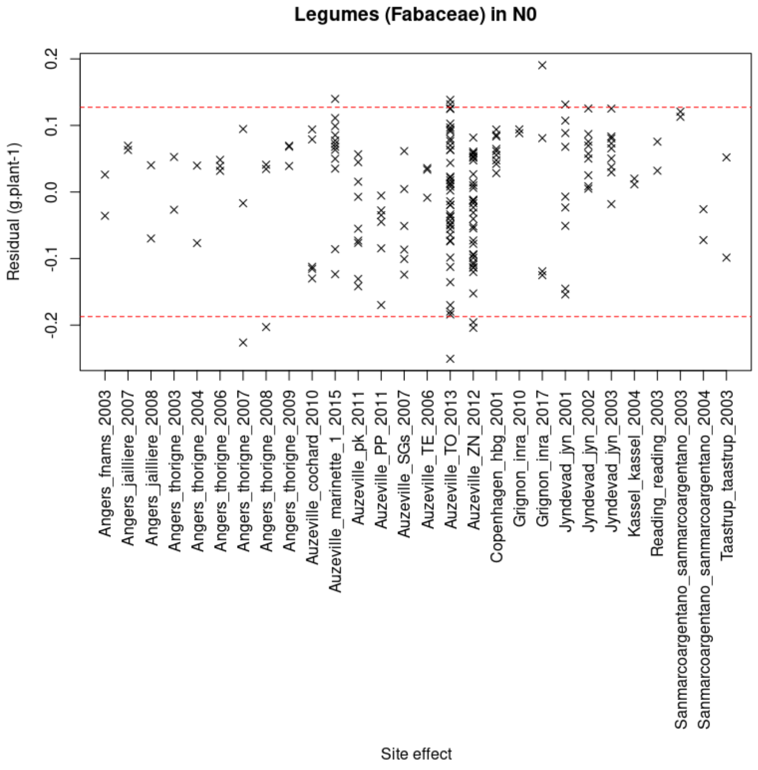


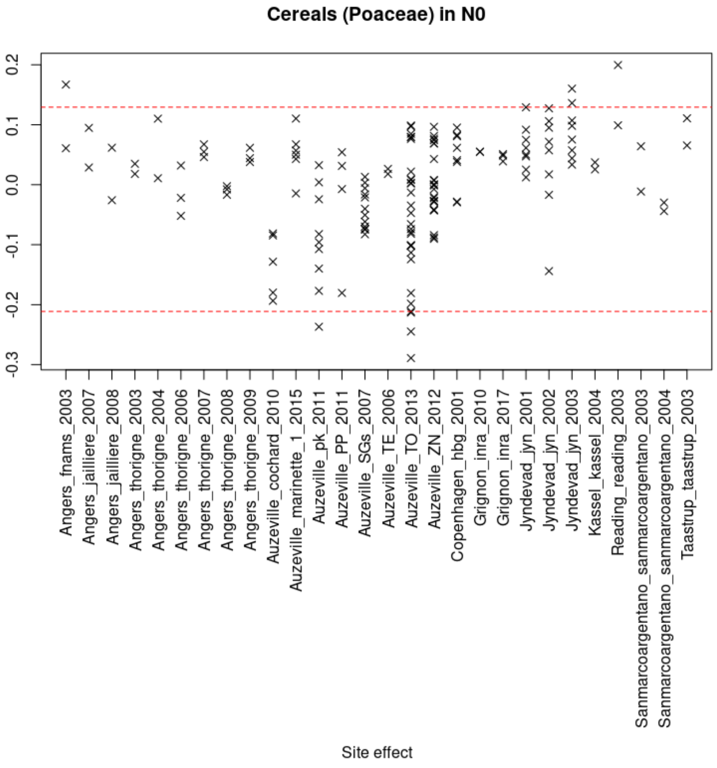


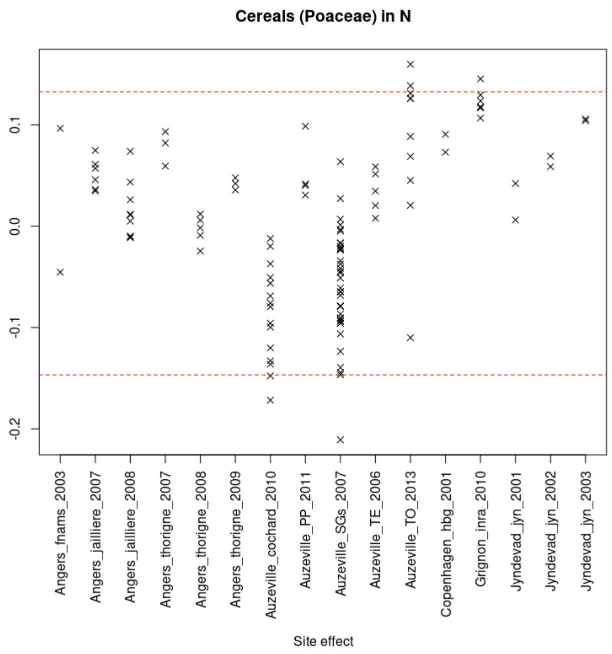
